# 1.7 GHz long-term evolution radiofrequency electromagnetic field with efficient thermal control has no effect on the proliferation of different human cell types

**DOI:** 10.1101/2023.09.25.559414

**Authors:** Jaeseong Goh, Dongwha Suh, Sangbong Jeon, Youngseung Lee, Nam Kim, Kiwon Song

## Abstract

Long-term evolution (LTE) radiofrequency electromagnetic field (RF-EMF) is widely used in communication technologies. As a result, the influence of RF-EMF on biological systems is a major public concern, and its physiological effects remain controversial. In our previous study, we showed that continuous exposure of various human cell types to 1.7 GHz LTE RF-EMF at specific absorption rate (SAR) of 2 W/Kg for 72 h can induce cellular senescence. To understand the precise cellular effects of LTE RF-EMF, we elaborated the 1.7 GHz RF-EMF cell exposure system used in the previous study by replacing the RF signal generator and developing a software-based feedback system to improve the exposure power stability. This refinement of the 1.7 GHz LTE RF-EMF generator facilitated the automatic regulation of RF-EMF exposure, maintaining target power levels within a 3% range and a constant temperature even during the 72-h exposure period. With the improved experimental setup, we examined the effect of continuous exposure to 1.7 GHz LTE RF-EMF at up to SAR of 8 W/Kg of adipose tissue-derived stem cells and Huh7, HeLa, and B103 cells. Surprisingly, the proliferation of all cell types, which displayed different growth rates, did not change significantly compared with that of the unexposed controls. However, when the thermal control system was turned off and the subsequent temperature increase induced by the RF-EMF was not controlled during continuous exposure to SAR of 8 W/Kg LTE RF-EMF, cellular proliferation increased by 35.2% at the maximum. These observations strongly suggest that the cellular effects attributed to 1.7 GHz LTE RF-EMF exposure were primarily due to the induced thermal changes, rather than the RF-EMF exposure itself.

## Introduction

Radiofrequency electromagnetic fields (RF-EMFs) are universally used in telecommunications and have become a daily necessity. In telecommunication technologies, 1.7 to 1.95 GHz long-term evolution (LTE) is widely used in 4^th^ generation mobile technologies [1, 2]. As LTE technologies have enabled the convergence of wired and wireless networks, such as GSM, LAN, and Bluetooth [3-5], LTE is currently the most widely adopted telecommunication technology. Despite the extensive daily use of these technologies, the physiological effects of LTE RF-EMFs on humans is not fully understood. However, these physiological effects are a major public health concern.

The International Commission on Non-Ionizing Radiation Protection (ICNIRP) defines specific absorption rate (SAR) of 2 W/kg as the safety limit for mobile device emissions; however, this limit remains controversial. Several studies have revealed the adverse effects, such as induction of DNA breakage and oxidative stress, of 1.8 GHz RF-EMF in human and mouse cells. Intermittent exposure (5 min on/10 min off) to SAR of 2 W/Kg 1.8 GHz RF-EMF for 24 h induced DNA single- and double-strand breaks in human fibroblasts and transformed GFSH-R17 rat granulosa cells [6]. In addition, 1.8 GHz RF-EMF exposure led to oxidative DNA damage (SAR of 4 W/Kg for 24 h) in mouse spermatocyte-derived GC-2 cells [7]. Xu et al. revealed that oxidative stress induced by exposure to SAR of 2 W/Kg 1.8 GHz RF-EMF might damage mitochondrial DNA, resulting in neurotoxicity [8]. However, these researchers also demonstrated that exposure to RF-EMF at SAR of 4 W/Kg did not elicit DNA damage [9]. Other studies have reported that RF-EMFs have no effects on mitochondrial function and did not induce apoptosis or chromosomal alterations; exposure to 1.95 GHz RF-EMF ranging from 0 to 4 W/Kg for up to 66 h did not induce apoptosis, oxidative stress, or DNA damage in human hematopoietic stem cells or the human leukemia cell line, HL-60 [10]. Intermittent exposure (5 min on/30 min off) to SAR of 1.5 W/Kg 1.71 GHz RF-EMF did not induce any significant cellular dysfunction in mouse embryonic stem cell-derived neural progenitor cells [11]. In addition, exposure to 900 MHz at 40 V/m for 1 h had no significant effect on the viability of human epidermal keratinocytes [12]. Exposure to 27.1 MHz RF-EMF did not influence the viability of the human keratinocyte cell line, HaCaT [13]. Thus, the effect of RF-EMF on cellular physiology remains controversial, and this uncertainty is another reason for public fear.

To understand the effects of RF-EMF in model organisms, the US National Toxicology Program (NTP) and the Ramazzini Institute in Italy conducted a carcinogenicity study of base-station exposure in mice and rats for more than 2 years. According to their findings, 900 MHz RF-EMF might induce cancer [14, 15]. However, the ICNIRP highlighted substantial limitations of the statistical analyses and stated that these loopholes preclude the conclusions drawn concerning RF-EMF and its carcinogenesis [16]. Owing to this controversy, continuous follow-up studies are required to identify the physiological changes induced by RF-EMF at both the cell and organism levels.

In our previous study, continuous exposure to SAR of 2 W/Kg 1.7 GHz LTE RF-EMF for 72 h inhibited the proliferation of various human cells by inducing cell senescence [17]. Using the same LTE RF-EMF generator employed by Choi et al. [17], we observed that exposure to SAR of 0.4 W/Kg 1.7 GHz LTE RF-EMF for 24 h activated cell proliferation while exposure to SAR of 4 W/Kg decreased the proliferation of human ASCs and Huh7 hepatocarcinoma cells (Supplementary Fig. S1 A and B). In this study, we aimed to understand the precise physiological effect of LTE RF-EMF on the proliferation of human cell types and elaborated the 1.7 GHz LTE RF-EMF cell exposure system previously used to eliminate the thermal effect. Using this refined RF-EMF exposure system, we investigated the effect of 1.7 GHz LTE RF-EMF on the proliferation of various mammalian cell types with different growth rates.

## Materials and Methods

### Sources of cells and culture

Human adenocarcinoma HeLa cells were purchased from the American Type Culture Collection, and human hepatocellular carcinoma Huh7 cells were purchased from the Korean Cell Line Bank. Human ASCs were purchased from Thermo Fisher Scientific (Waltham, MA, USA). B103 rat neuroblastoma cells were a gift from Dr. Inhee Mook-Jung (Seoul National University College of Medicine, Seoul, Korea).

HeLa, B103, and Huh7 cells were cultured in high glucose-containing Dulbecco’s modified Eagle’s medium (DMEM; Gibco) supplemented with 10% fetal bovine serum (FBS; Sigma-Aldrich, St. Louis, MO, USA) and 1% penicillin-streptomycin (Gibco). ASCs were grown in DMEM/F12 (Gibco) supplemented with 10% FBS and 1% penicillin-streptomycin. All cell types were cultured at 37 °C in a humidified atmosphere containing 5% CO_2_.

### Cell exposure to the LTE RF-EMF radiation system

The exposure system was preheated for a minimum of 30 min before 1.7 GHz LTE RF-EMF exposure. A total of 30 × 10^4^ ASCs and 20 × 10^4^ Huh7, HeLa, and B103 cells were seeded and incubated in a 100-mm dish for 16 h before RF-EMF exposure. For RF-EMF exposure, as previously described [17], 100-mm culture dishes were placed 13.6 cm from the conical antenna, which was located at the center of the exposure chamber. The cells in the dishes were then exposed to the RF-EMF of a single LTE signal at SAR values ranging from 0.4 to 8 W/Kg for the described duration. During the exposure, the temperature of the exposure chamber was maintained at 37 ± 0.5 ºC by circulating water within the chamber. The unexposed sham group was incubated under the same conditions without RF-EMF exposure. After RF-EMF exposure, the cells were counted using Cellometer Auto T4 (Nexcelom) and used for further assays.

### Cell viability assay

Cell viability was monitored using cell counting and MTT assays. After exposure to RF-EMF for the indicated time periods, the cells were harvested and counted using a Cellometer Auto T4 (Nexcelom). For the MTT assay, Huh7, HeLa, and B103 cells were incubated in 6 mL of medium containing 0.5% 3-(4,5-dimethylthiazol-2-yl)-2,5-diphenyltetrazolium bromide (MTT; Amresco Inc., OH, USA) at 37 °C for 90 min and ASCs for 3 h. The resulting formazan crystals were dissolved in 6 mL of dimethyl sulfoxide (DMSO), and the optical density was measured at 570 nm using an ELISA microplate reader (SpectraMax ABS, Molecular Device Co., CA, USA).

### Statistical analysis

All statistical analyses were performed using GraphPad Prism 9 (GraphPad Software Inc., San Diego, CA, USA). All data are presented as mean ± standard deviation (SD) of more than three independent experiments with statistical significance. P<0.05 (*), P<0.01 (**), and P<0.001 (***) were considered to indicate statistical significance while P>0.05 was considered to indicate statistical non-significance (ns).

## Results

### New RF-EMF signal generator and a software-based target power feedback system increase the stability of 1.7 GHz LTE RF-EMF signal generation

The LTE RF-EMF cell exposure system consists of an incubator, water circulator, signal generator, power meter, power amplifier, and control computer. A JS-CO2-AT-750 incubator (John Sam Corp.) was used to maintain a controlled environment for the cell culture, and a C-WBL water circulator (Chang-shin Science) was used to provide cooling and eliminate any thermal effects during exposure. To refine the system, a new signal generator, E4438C (Keysight Technologies), was employed to generate the LTE signal. The generated LTE signal was amplified using a customized power amplifier developed to ensure a maximum output power of 60 W, and an E4418B power meter (Keysight Technologies) was employed to measure the amplifier output powers (Fig. 1A).

**Fig. 1.**
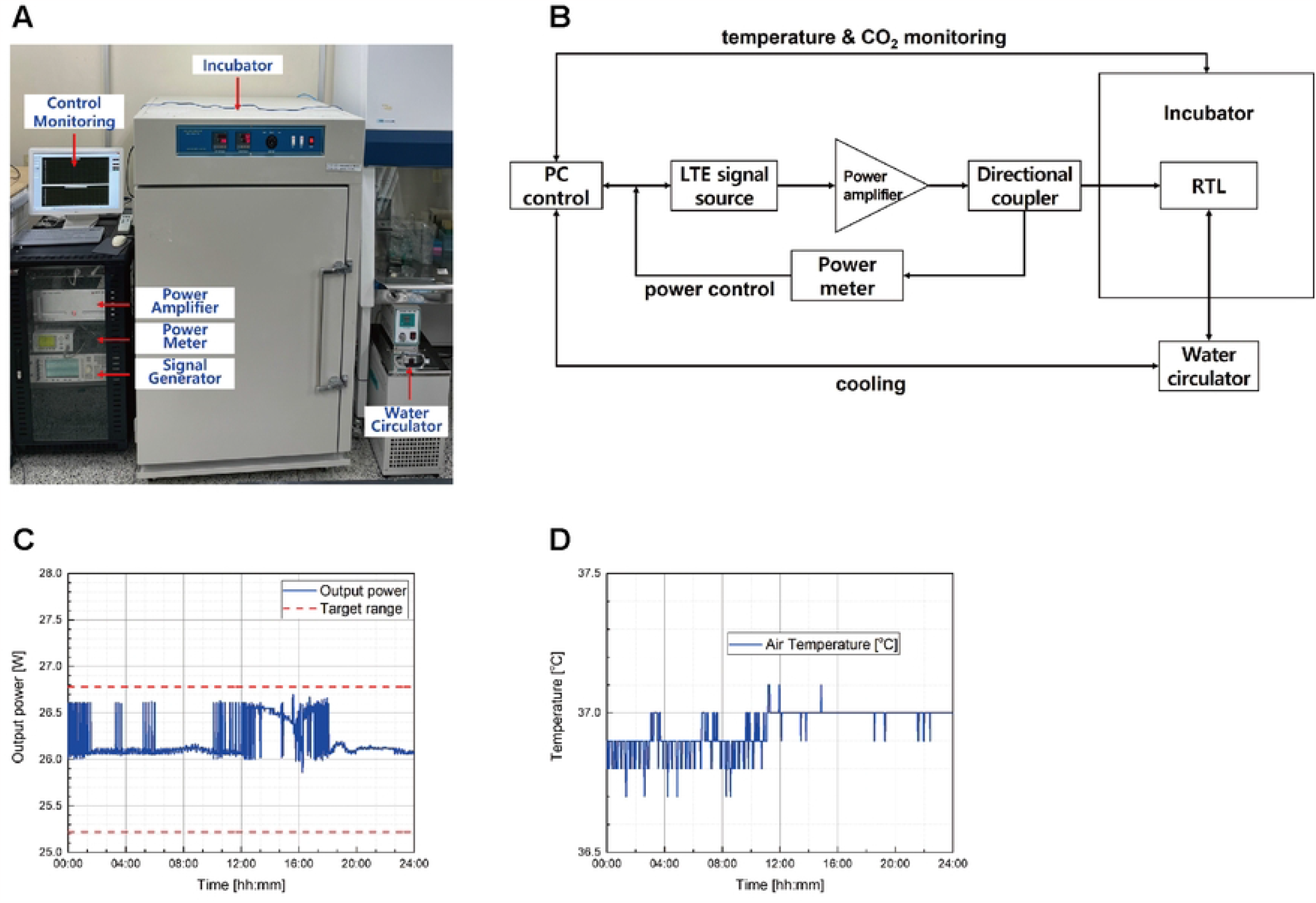
Elaborate 1.7 GHz LTE RF-EMF cell exposure system used in this study. (A) Image of the 1.7 GHz RF-EMF exposure device. (B) Block diagram of the 1.7 GHz LTE RF-EMF exposure system. (C) The output power was monitored for 24 h. (D) The air temperature of the incubator was monitored for 24 h.

To continuously monitor the power generated by E4418B, a customized control software was developed and installed on a PC to display the power measurement results (Fig 1B). During the exposure, the PC continuously recorded all experimental data and controlled all feedback flows to regulate the desired conditions and settings in real time. All exposure conditions, such as frequency, duration, and signal level, can be defined using the main software. The LTE signal was amplified to a desired level using a power amplifier and injected into a radial transmission line. The power measured through a directional coupler and a power meter functions as a feedback loop connected to an LTE signal generator, which maintained the power within ± 3% of the target output power level. This feedback scheme was implemented to regulate the one-minute average output power, which is crucial for maintaining the required SAR values in a stable manner.

To check the stability of the output power through power control, we set the target power to 26 W and the target range to ± 3%, and then monitored the output for 24 h. An example of power monitoring is presented in Fig 1C, which shows that the output power was well controlled within the target range (Fig. 1C). The mean output was 26.12 W, and the standard deviation was 0.76%. The air temperature in the incubator was also well controlled, with a mean of 36.95 °C (Fig. 1D).

### 1.7 GHz LTE RF-EMF generated with the refined system does not affect the proliferation of various mammalian cells

We previously reported that exposure to 1.7 GHz LTE RF-EMF at SAR of 2 W/Kg for 72 h induced senescence in ASCs and Huh7 cells [17]. In addition, the number of ASCs and Huh7 cells decreased by 7% and 20%, respectively, following exposure to the same system at SAR of 4 W/Kg for 24 h (Supplementary Fig. S1 B). Using the same experimental setup, when ASCs and Huh7 cells were exposed to SAR of 0.4 W/Kg for 24 h and incubated for an additional 48 h, the numbers of ASCs and Huh7 cells increased by 27% and 37%, respectively, compared with that of the unexposed control (Supplementary Fig. S1 A). However, with the new elaborate RF-EMF exposure equipment setup, under the same exposure conditions used in the previous setup (24 h of exposure followed by 48 h of incubation), neither ASCs nor Huh7 cells showed any significant change in cell number after exposure to SAR of 0.4 W/Kg and 4 W/Kg, compared to the unexposed controls (Fig. 2A).

**Fig. 2.**
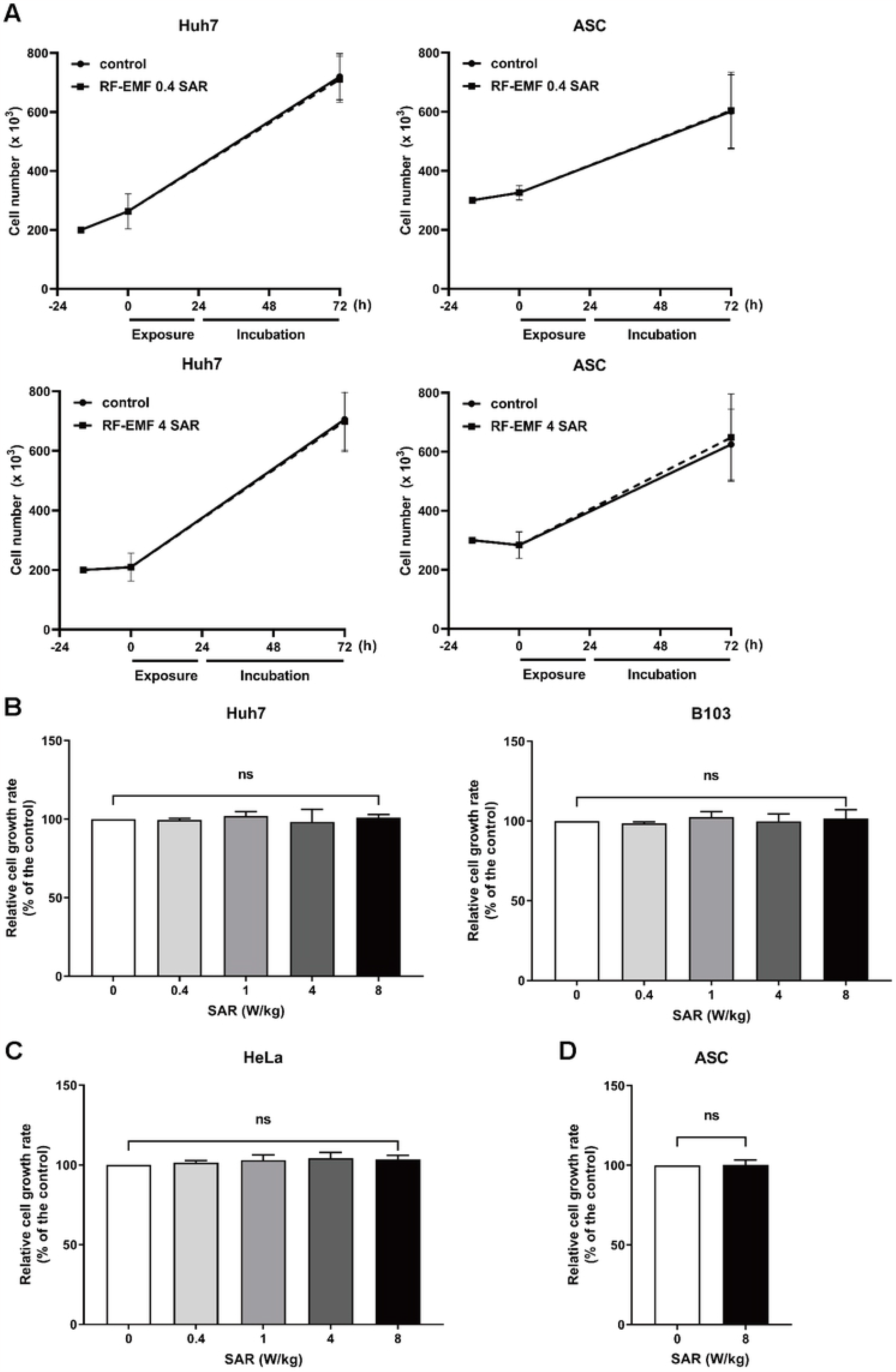
Exposure to 1.7 GHz LTE RF-EMF generated in this refined system did not affect cellular proliferation. The same number of cells was seeded and incubated for 16 h. (A) Huh7 cells and ASCs were exposed to the SAR of 0.4 W/Kg or 4 W/Kg RF-EMF for 24 h and further incubated for 48 h without RF-EMF exposure. After incubation, the cells were counted using a cell counter. (B) Huh7, HeLa, and (C) B103 cells were exposed to RF-EMF for 72 h at the indicated SAR values. After exposure, cell viability was assessed using the MTT assay. (D) ASCs were exposed to the SAR of 8 W/Kg RF-EMF for 72 h, and cell viability was assessed using the MTT assay. Zero SAR: Unexposed controls. More than three independent replicates were tested in all experiments, and the data are presented as mean ± SD. ns represents non-significant.

To assess the effects of exposure to the 1.7 GHz LTE RF-EMF generated using this refined system on cell proliferation, we evaluated cell viability by extending the exposure time to 72 h, with the exposure intensity ranging from SAR of 0.4 W/Kg to 8 W/Kg, and performing MTT assays. If the exposure to RF-EMF positively or negatively affects cell proliferation, the effect would be more obvious in fast-growing cells. Thus, we used the fast-growing human cancer cells, Huh7 and HeLa, for the exposure and viability assays. We also exposed B103 rat neuroblastoma cells, which grow faster than human cancer cells, to LTE RF-EMF. Only the effect of RF-EMF exposure at SAR of 8 W/Kg was monitored in ASCs; this is because stem cells, including ASCs, usually proliferate slowly, and exposure effects would not be obvious at low intensity. The viability of Huh7, HeLa, and B103 cells exposed to 1.7 GHz LTE RF-EMF was not significantly different from that of the unexposed controls at all tested SAR values (Fig. 2B and C). Correspondingly, at SAR of 8 W/Kg, no significant difference was noted in the viability of exposed ASCs compared with that of the unexposed control (Fig. 2D).

### Exposure to 1.7 GHz LTE RF-EMF activates cell proliferation without the temperature control system

No cellular effect of exposure to 1.7 GHz LTE RF-EMF generated by our new elaborate system was observed, although pro- and anti-proliferative effects of 1.7 GHz LTE RF-EMF were observed, depending on the SAR values, in our previous generator setup. Thus, we speculate that the temperature elevation caused by RF-EMF generation in the previous setup might be responsible for the observed cellular effects of the RF-EMF. To confirm this speculation, we monitored the chamber temperature and evaluated the cell viability without turning on the water circulation system. When the water circulator was turned off, the chamber temperature increased by approximately 1.7 °C compared to that obtained when the circulator was turned on (Fig. 3A). When B103 and HeLa cells and ASCs were exposed to 1.7 GHz LTE RF-EMF at the SAR of 8 W/Kg without maintaining the temperature by water circulation, we observed a significant acceleration in cell proliferation depending on the cell growth rate: 11% for ASCs, 24.3% for HeLa, and 35.2% for B103 (Fig 3B).

**Fig. 3.**
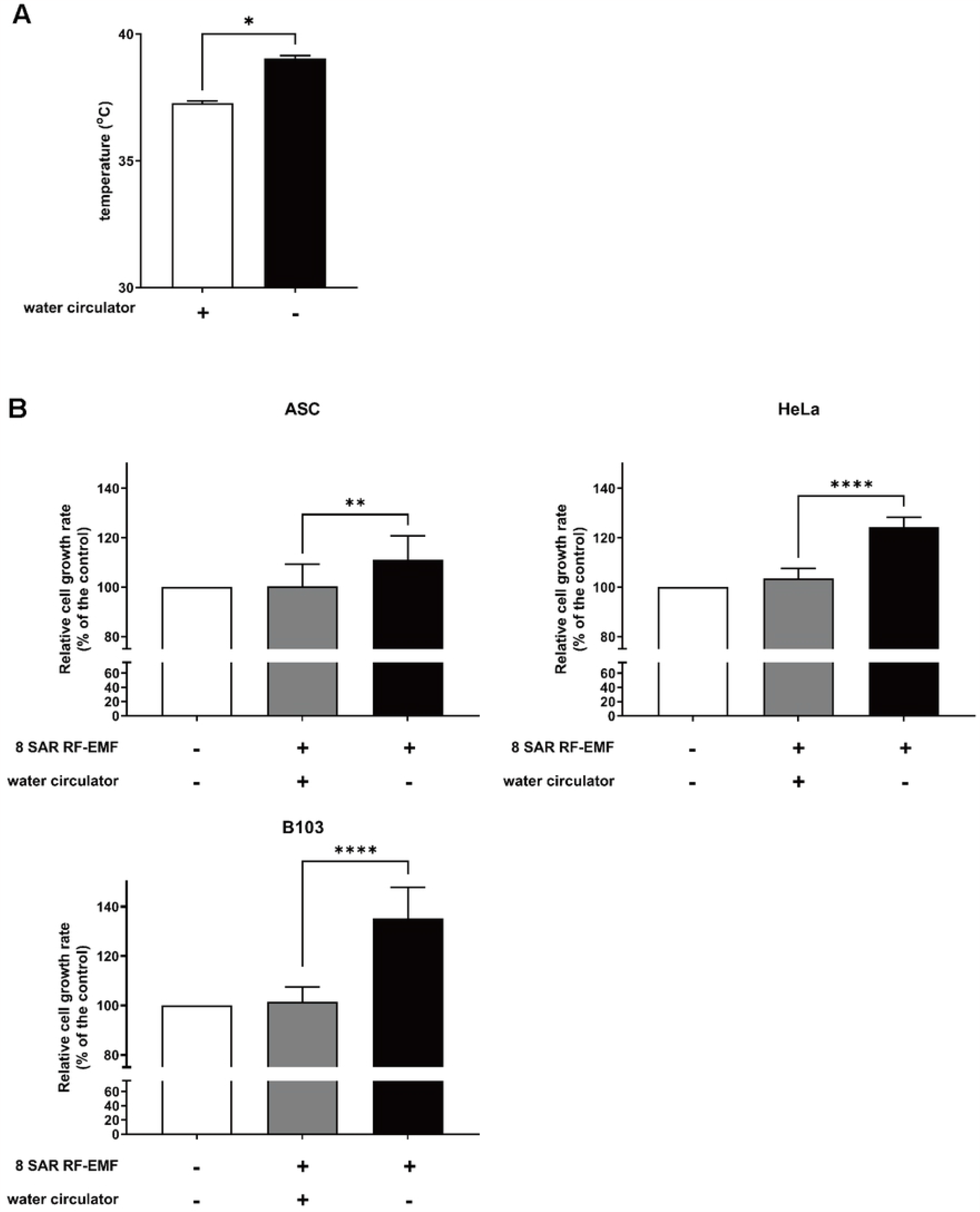
Exposure to 1.7 GHz LTE RF-EMF without heat control activated cell proliferation due to thermal effects. (A) During exposure to the SAR of 8 W/Kg 1.7 GHz LTE RF-EMF, the temperature within the exposure chamber was recorded every minute for 72 h, and the average temperature was calculated. The temperature was measured when the water circulator was turned on or off. (B) ASCs and HeLa and B103 cells were exposed to the SAR of 8 W/Kg 1.7 GHz RF-EMF for 72 h, while the water circulator was turned on or off. Cell proliferation was measured using the MTT assay. More than three independent replicates were tested in all experiments, and the data are presented as mean ± SD. P < 0.0001 (****), P < 0.01 (**).

## Discussion

In this study, 1.7 GHz LTE RF-EMF from a stabilized generating system with minimal thermal effect was demonstrated to have no effect on the proliferation of various mammalian cell lines with different growth rates. However, when the temperature was not adequately controlled, cell proliferation was promoted upon exposure to RF-EMF. These observations strongly suggest that temperature is a major underlying factor for the previously reported cellular effects of 1.7 GHz LTE RF-EMF.

Temperature is a well-established factor that influences various cellular processes, including metabolic activity and enzymatic reactions. Various biological species exhibit a temperature-dependent increase in cell proliferation within specific temperature ranges [18]. For instance, HeLa cells exhibit a temperature-dependent increase in growth rate between 33 and 38 ºC [19]. Conversely, temperatures outside a certain range can have adverse effects on cell growth, inducing cold-or heat-shock responses. Heat shock stress induces the unfolding of intracellular proteins, as well as cytoskeleton and cell membrane damage. Consequently, accumulation of heat stress can lead to cell cycle arrest or cell death [20]. Furthermore, heat stress-induced alterations in the mitochondrial antioxidant system can result in increased ROS generation [21].

As RF-EMF leads to temperature elevation, investigation of the direct effects of RF-EMF on cells and biological organisms requires precise temperature control and monitoring. In our case, switching the RF-EMF generator and implementing a feedback monitoring system enabled more consistent generation of LTE RF-EMF with stable intensity. As a result, stabilization of the intensity contributes to an increase in thermal stability. Using this improved RF-EMF exposure system, we verified that the anti- and pro-proliferative effects observed and reported in our previous studies were mainly due to thermal effects. Future research groups that assess the cellular or physiological effects of RF-EMF should compare their results with a more elaborate RF-EMF system that has better thermal control. We also expect that the influence of 1.7 GHz LTE RF-EMF exposure, at least up to SAR of 8 W/Kg, on biological organisms would be minimal, as biological organisms usually have better thermal control systems for the maintenance of homeostasis. Nonetheless, our results should be confined to 1.7 GHz LTE RF-EMF, and RF with different frequencies might have different physiological outcomes.

## Conclusion

To understand the precise cellular effect of 1.7 GHz LTE RF-EMF, we developed an RF-EMF cell exposure system with an improved RF signal generator and control software. With a refined RF-EMF exposure system, we could maintain a consistent target power during the 72-h exposure period with minimal thermal effects. With this refined experimental setup, exposure to 1.7 GHz LTE RF-EMF at the SAR ranging from 0.4 W/Kg to 8 W/Kg was not found to affect the proliferation of various human cells with different growth rates. Before upgrading the exposure system, we observed that the exposure of human cells to 1.7 GHz RF-EMF increased or decreased cell proliferation, depending on the SAR values. In addition, we verified that exposure to 1.7 GHz RF-EMF with this refined system affected cell proliferation when heat was not properly controlled. Altogether, these results suggest that exposure to 1.7 GHz LTE RF-EMF does not directly influence cell proliferation and that the RF-EMF effects might be associated with thermal effects.

## Abbreviations used

LTE: long-term evolution
RF-EMF: radiofrequency-electromagnetic field
ASC: adipose tissue-derived stem cell
SAR: specific absorption rate

## Acknowledgments

This study was supported by the ICT R&D program of MSIT/IITP [2019-0-00102, A Study on Public Health and Safety in a Complex EMF Environment]. D. Suh and J. Goh were partially supported by the BK21 PLUS and BK21 FOUR programs, respectively.

## Supporting information

**Supplementary Fig. S1. Exposure to 1.7 GHz LTE RF-EMF with the previous generating system induced either a positive or negative effect on cellular growth depending on the SAR values**. Equal amounts of ASCs and Huh7 cells were seeded and incubated for 16 h. The cells were then exposed to the SAR of (A) 0.4 W/Kg or (B) 4 W/Kg 1.7 GHz LTE RF-EMF for 24 h, and further incubated for 48 h without RF-EMF exposure. After incubation, the cells were counted using a cell counter and plotted. Three independent experiments were performed, and the cell number is presented as mean ± SD. P < 0.001 (***), P < 0.05 (*).

## Notes

### Competing Interest Statement

The authors have declared no competing interest.

